# A specific relationship between musical sophistication and auditory working memory

**DOI:** 10.1101/2021.07.08.451659

**Authors:** Meher Lad, Alexander J. Billig, Sukhbinder Kumar, Timothy D. Griffiths

## Abstract

Musical engagement may be associated with better listening skills, such as the perception of and working memory for notes, in addition to the appreciation of musical rules. The nature and extent of this association is controversial. In this study we assessed the relationship between musical engagement and both sound perception and working memory.

We developed a task to measure auditory perception and working memory for sound using a behavioural measure for both, *precision*. We measured the correlation between these tasks and musical sophistication based on a validated measure (the Goldsmiths Musical Sophistication Index) that can be applied to populations of both musicians and non-musicians. The data show that musical sophistication accounts for 21% of the variance in the precision of working memory for frequency in an analysis that accounts for age and non-verbal intelligence. Musical sophistication was not significantly associated with the precision of working memory for amplitude modulation rate or with the precision of perception of either acoustic feature.

The work supports a specific association between musical sophistication and working memory for sound frequency.

## Introduction

Musical engagement involves processing at multiple sensory, perceptual and cognitive levels. In the pitch domain, it requires an analysis of notes with different sensory properties (frequency structure, temporal envelope and fine-temporal structure) that are associated with different perceptual pitch values. Notes form phrases that are held in mind over short periods of time and compared, a process that requires auditory working memory. The structure of phrases is determined by musical rules that can be learned during musical engagement either implicitly by exposure or explicitly by instruction. Additionally, instrumental performance requires learned sensorimotor coordination.

We consider here the extent to which musical engagement, of any kind, is associated with improved listening skills: sound perception and working memory. Previous studies have examined fundamental perception, often based on comparison between groups of musicians and controls. Musicians have been shown to have significantly lower thresholds for frequency discrimination than non-musicians (Kishon-Rabin et al., 2001): classical musicians had an advantage over those trained in contemporary music. Additionally, musicians such as guitarists and flautists who have to adjust the pitch of instruments discriminate finer frequency differences than those playing fixed-pitch instruments like piano (Micheyl et al., 2006). However, the effect of musicianship on frequency discrimination has not been consistently demonstrated (Moore et al., 2019).

It has been argued that the effect of musical sophistication on perceptual tasks is determined by cognitive abilities (Ahissar et al., 2009). Auditory working memory for tones is an explicit cognitive task that has been examined in a number of studies in which improved performance is shown in groups of musicians compared to non-musicians (Ding et al., 2018; Talamini et al., 2021; Williamson et al., 2010). Musicians have also been shown to have improved sustained attention (Carey et al., 2015) and attention to pitch direction (Ouimet et al., 2012), as well as better general auditory cognition in terms of phonological working memory and speech-in-noise perception (Parbery-Clark et al., 2011, 2009).

Previous studies of the relationship between musicality and perceptual and cognitive abilities have used perceptual and cognitive tasks that differ markedly from each other (Carey et al., 2015). There is a need in this area for perceptual and cognitive tasks that are more closely aligned in methodology and output metrics to facilitate comparison of task effects. In this study, we used a measure of precision (inverse of the standard deviation) both for perception and working memory. This measure is related to errors in matching sounds to a target that are immediately adjacent in time (perceptual precision) or separated in time by seconds (working memory precision). Previous studies of auditory working memory are consistent with the distribution of errors reflecting the cognitive resource allocated to working memory for the sound feature (Kumar et al., 2013). We have adapted the delayed matching design used for working memory error measurement to allow similar measures of perceptual and working memory precision.

Moreover, previous studies have often used categorical designs based on the comparison of groups of musicians and non-musicians. In this study we took a correlational approach based on a continuous measure of musical experience that can be applied to any subject without imposing an arbitrary boundary between musicians or non-musicians. The approach is potentially more powerful and recognizes the fact that we all have musical experience, even if we are not formally trained. The measure is the Goldsmiths Musical Sophistication Index (Gold-MSI) (Müllensiefen et al., 2014), a validated tool to assess self-reported musical skills and behaviours that captures the variety of musical behaviours in society.

We demonstrate correlation of musical sophistication with working memory precision for frequency, but not with working memory precision for amplitude modulation rate, nor with perceptual precision for either sound feature. The work defines a specific listening skill related to musical sophistication.

## Results

Experiment 1 (Fig 1) measured auditory working memory precision for tone frequency and for amplitude modulation (AM) rate of sinusoidally modulated noise. It also used a control task based on working memory precision for colour and flash rate of visual stimuli. In both paradigms subjects had to manually adjust a test stimulus to a target stimulus heard before a delay.

**Figure 1.**
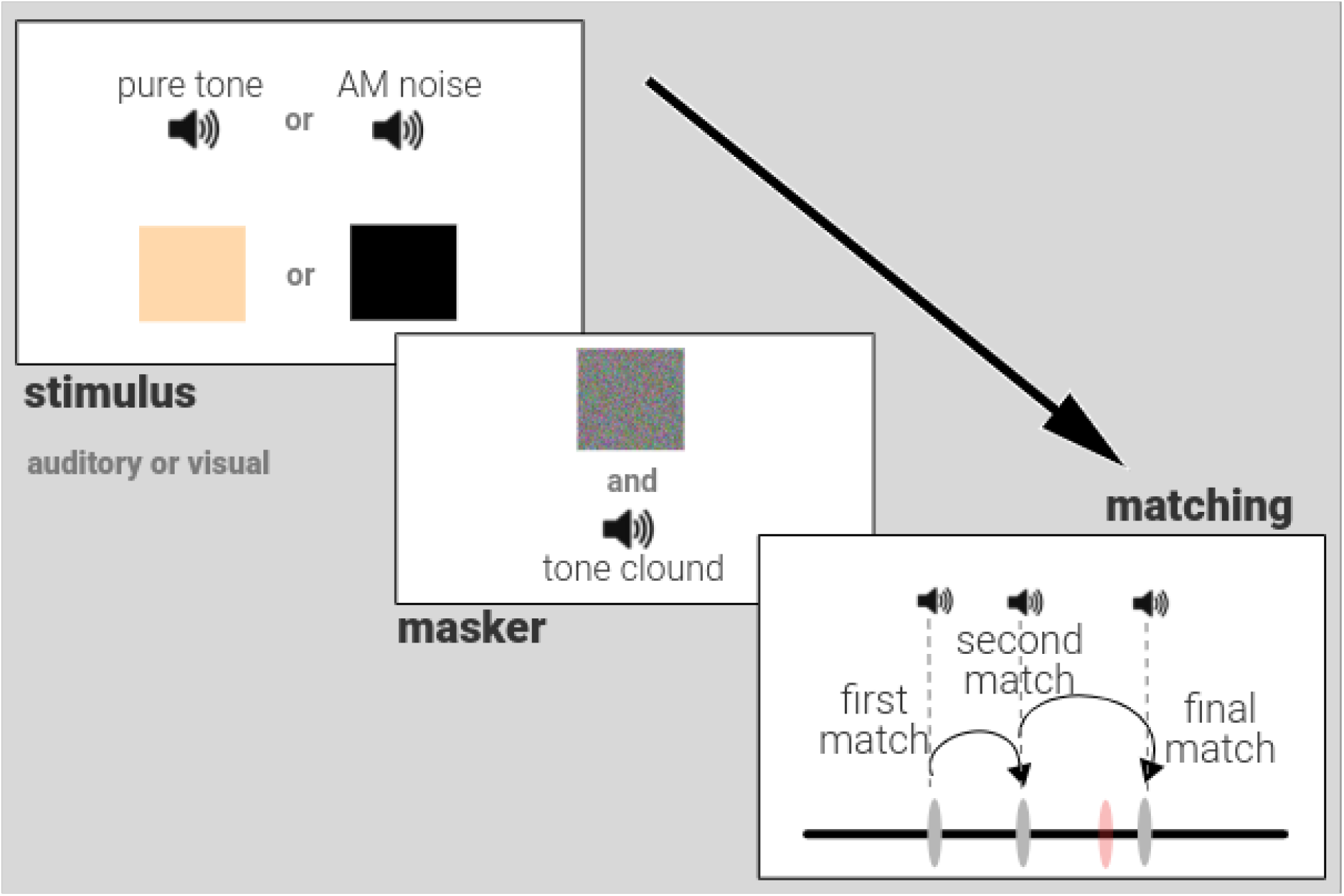
Description. *Working Memory Precision Experiment*. After an auditory (pure tone or amplitude modulated noise) or a visual (colour or flashing box) stimulus is presented for 1 second, a brief masker (visual and auditory) is presented for 0.5 seconds. After a subsequent delay of 2 to 4 seconds, participants can match to the original stimulus using a horizontal scale on the screen. The scale is linked to the parameter of interest (frequency for pure tone or AM rate) that generates the original stimulus on a given trial and participants can explore the parameter space to ‘find’ the stimulus. The figure shows an auditory matching trial where the participant’s ‘final match’ (dark grey marker on the scale) is shown in comparison to where the original stimulus (orange marker on the scale) is actually located. In this example, the participant first clicked on the scale to make a ‘first match’ (which produced a sound linked to the parameter at that location), then a ‘second match’ and then a ‘final match’. The discrepancy between the ‘final match’ location parameter and that of the original stimulus gives an ‘error’ for each trial that can be used to calculate the auditory working memory ‘precision’, the inverse of the standard deviation of errors from a trial target, for all auditory trials.

Age and non-verbal reasoning scores were used as regressors of no interest for all analyses. There was a positive correlation between Gold-MSI scores and precision of auditory working memory for frequency, *r*(98) = 0.46, *p* < 0.001 (Fig 2B). There was no statistically significant correlation between Gold-MSI and precision of auditory working memory for AM rate, *r*(98) = 0.10, *p* = 0.334, visual working memory for colour precision, *r*(50) = −0.10, *p* = 0.802, or flash rate precision, *r*(50) = −0.09, *p* = 0.496. There was a significant difference between the correlation coefficients of auditory working memory for frequency and amplitude modulation rate precision, and the Gold-MSI, *p* < 0.001 (bootstrapped), after permutation testing.

**Figure 2.**
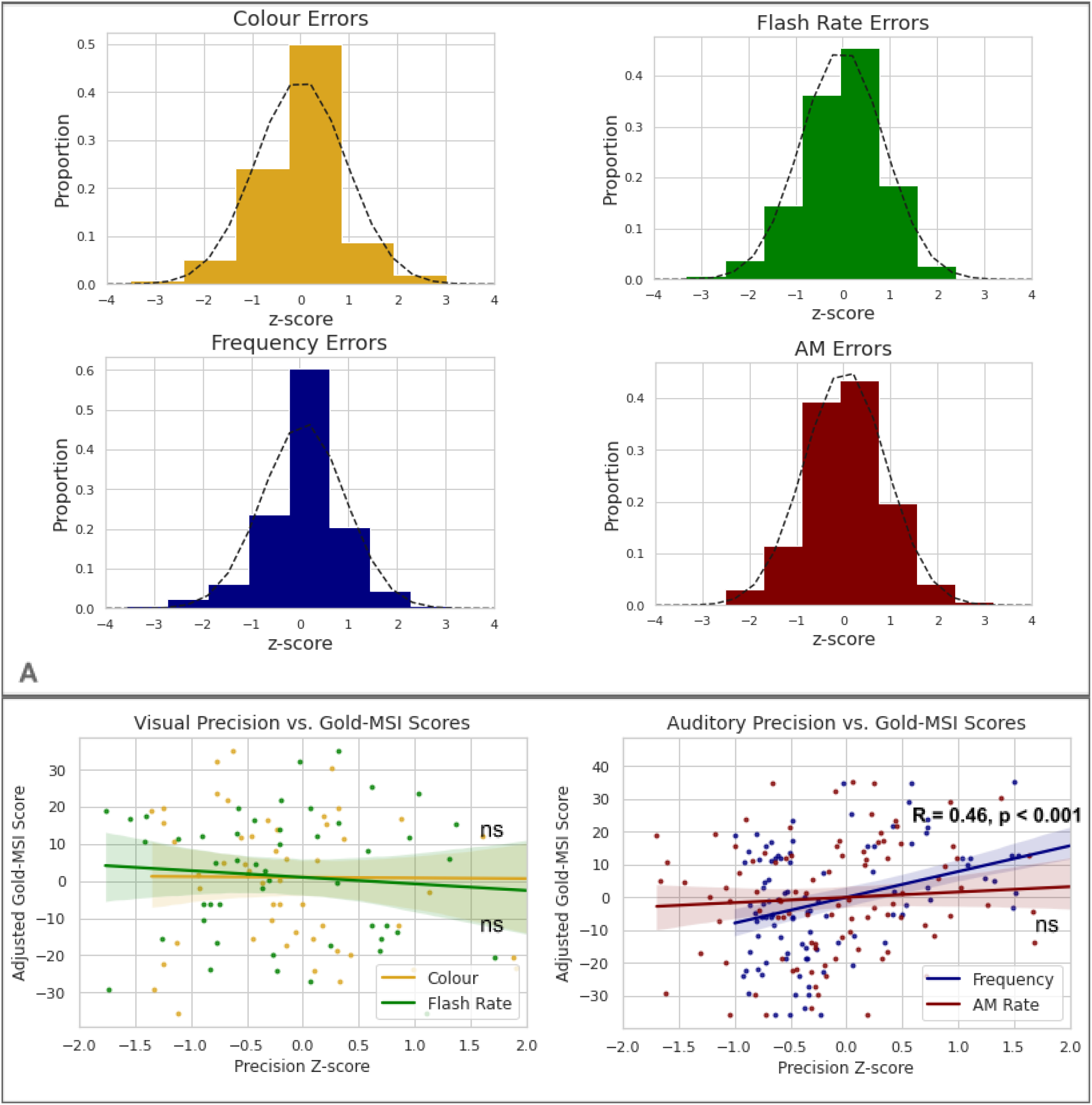
Description. 2A) The distributions for all domains of working memory tested in our experiment are shown. All participants’ performances were Gaussian. Z-scores are used for errors on the x-axis for the purposes of illustration. 2B) Scatter plots showing overall working memory precision for visual (top row) and auditory (middle row) stimuli as a function of transformed scores on the Gold-MSI questionnaire. Performance with visual stimuli was not significantly correlated with Gold-MSI scores. For auditory stimuli, working memory precision for frequency (represented by the dark blue line) was significantly correlated with Gold-MSI scores, but working memory precision for AM rate (represented by the red line) was not. ns - not significant

Experiment 2 (Fig 3) measured the precision of auditory working memory for frequency and AM rate in addition to the *perceptual* precision of frequency and AM rate. The aim was to identify whether musical sophistication was associated specifically with the more cognitive task that required holding a sound in mind for several seconds, or if the association extended to processing of sound over shorter timescales. Additionally, the experiment allowed a further test of the correlation between musical sophistication and auditory working memory for frequency in an independent cohort.

**Figure 3.**
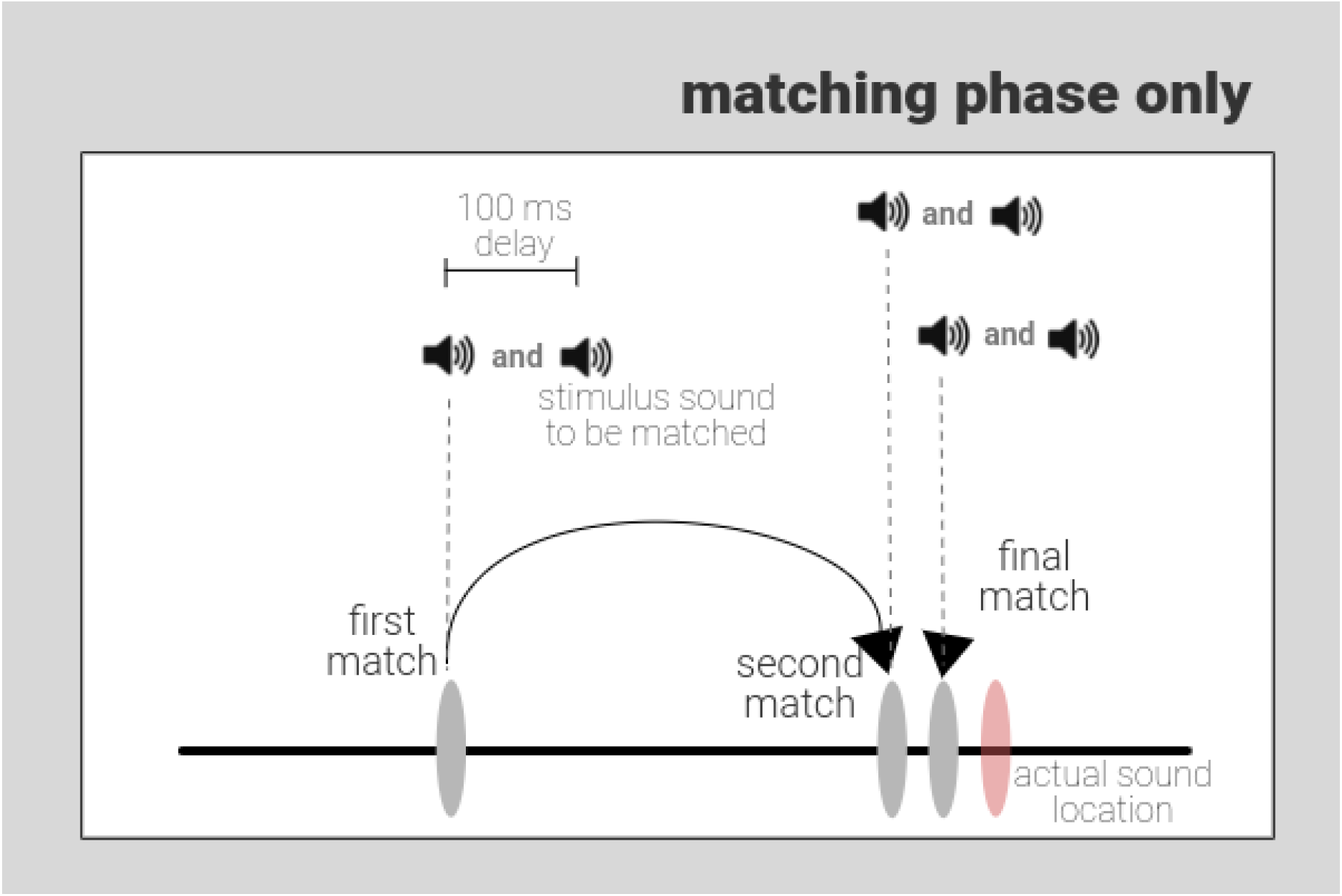
Description. *Perceptual Precision Experiment*. In comparison to Experiment 1, there is only a matching phase. The stimulus (auditory trial shown here) to be matched is presented 100 milliseconds after a match is made on the scale and so each click on the scale produces two sounds in a sequence. The first is the sound at the parameter location of the click and the second is the sound to match to. Participants are asked to click multiple times on the scale (hence, ‘first match’, ‘second match’ and ‘final match’) until they find a point where the two sounds in succession sound the same. In this auditory trial, the light red ellipse indicates the true location of the sound parameter to be matched.

Neither auditory perceptual precision for frequency, *r*(44) = 0.23, *p* = 0.121, nor perceptual precision for AM rate, *r*(44) = 0.27, *p* = 0.061 correlated with Gold-MSI scores (Fig 4B). The correlation between the Gold-MSI and precision of working memory for frequency was significantly greater than that between Gold-MSI and precision of frequency *perception*, *p* < 0.001 (bootstrapped), after permutation testing. Furthermore, the correlation for precision of working memory was significant even after adjusting for frequency perceptual precision (in addition to age and non-verbal reasoning), *r*(43) = 0.37, *p* = 0.009.

**Figure 4.**
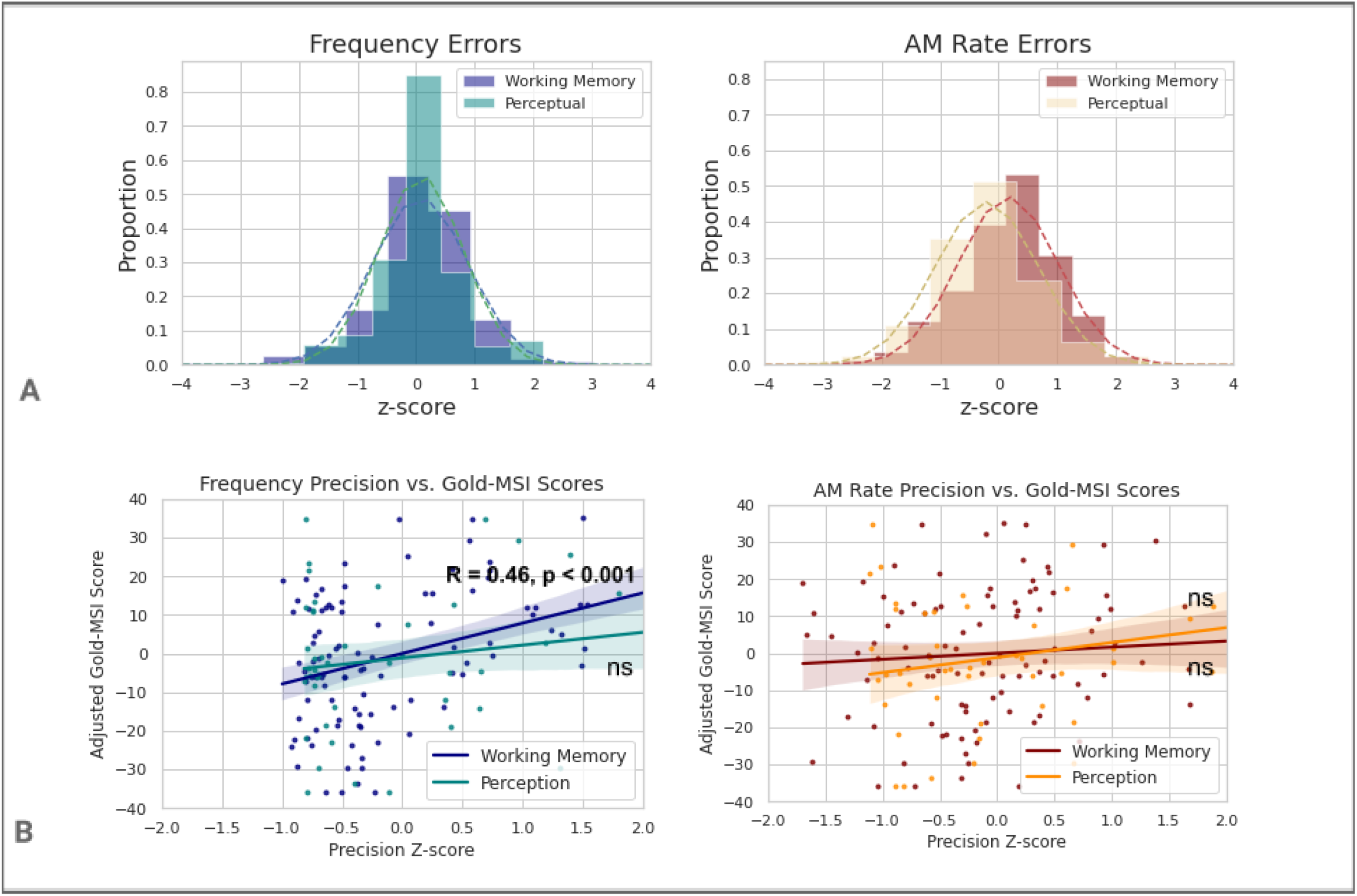
Description. 4A) The distributions for all auditory domains of working memory and perception tested in our experiment are shown (top row). All participants’ performances were Gaussian but the distribution of perceptual errors has a smaller degree of variability than working memory errors. 4B) Scatter plots showing overall auditory precision for frequency (left) and amplitude modulation rate (middle row) as a function of scores on the Gold-MSI questionnaire. For tone frequency, precision of working memory (represented by the dark blue line) but not perception (represented by the teal coloured line) was significantly correlated with Gold-MSI scores. For AM rate, neither perceptual nor working memory precision was significantly correlated with Gold-MSI scores. ns - not significant

## Discussion

This study demonstrates a significant correlation between the precision for frequency working memory and musical sophistication. The correlation is specific to the auditory domain and is not explained by age or non-verbal intelligence. The correlation is present for the precision of frequency working memory but not the precision of AM working memory.

### Musical sophistication is related to frequency but not AM listening skills

Frequency is a determinant of pitch, a fundamental aspect of music. Changes in pitch form the basis of melody and combinations of pitch form the basis of harmony. These aspects of music are common to many musical cultures and styles. Our study tested participants using a frequency range between 440 and 880 Hz, corresponding to the fundamental frequencies of notes commonly encountered in western music. The range of amplitude modulation rates used in our study (below 20 Hz) is not associated with pitch. It is within the range that tremolo can be added to sung and instrumental notes, for which note-by-note comparison or comparison over phrases is not as critical to the musical experience. Our findings are congruent with previous studies showing a particular advantage in working memory for tonal over verbal and visuospatial stimuli in musicians: see (Talamini et al., 2017) for meta-analysis.

### Musical sophistication is related to working memory precision but not perceptual precision

Previous work suggested that musicians have lower frequency discrimination thresholds than non-musicians (Carey et al., 2015; Kishon-Rabin et al., 2001). Other work suggests that working memory may drive these effects in certain circumstances (Ahissar et al., 2009; Zhang et al., 2016). For example, when the standard is varied on a trial-by-trial basis, working memory might influence performance. Working memory resources help create a ‘perceptual anchor’ when the representations can be easily degraded or interfered with (Nahum et al., 2010). This is in contrast to fixed standards that are often used to measure frequency difference limens. In this study we have used a stimulus based on a general precision score for standards that are varied in frequency for our perceptual task. However, we do not see any effect of musical sophistication on this perceptual precision. In contrast, we see a clear effect when explicit working-memory demands are imposed based on a longer delay and distractor. This dissociation between similar tasks that differ in working memory demands strongly supports a specific relationship between working memory for frequency and musical sophistication.

### Critical aspects of musical sophistication

Musical sophistication may include aural skills, receptive responses and the ability to make music (Hallam and Prince, 2003). The Gold-MSI captures this multifaceted nature of musical expertise and is shown to correlate with listening tests of musical ability in the form of melodic memory and musical beat perception (Müllensiefen et al., 2014). The key elements of the Gold-MSI include five domains which have been derived from factor analysis of a sample of around 150,000 participants. These domains include active engagement, perceptual abilities, musical training, singing abilities and emotional experience. This method is better suited to detect benefits to (or associations with) musicality offered by music communication, journalism or DJing, for example, where instrumental expertise or instruction may not have been attained. Further studies might address an association between specific domains and working memory for frequency using the full version of the Gold-MSI.

### Causal relationship between musical sophistication and listening skills?

We have demonstrated a specific correlation between musical sophistication and a listening skill that is not explained by a general effect of intelligence. We consider here the nature of possible underlying causal relationships between musical sophistication and listening skills. The current study, however, cannot provide a definitive answer about these.

Better innate general listening skills might allow better musical listening and the acquisition of musical sophistication. General natural listening and musical listening both require sustained attention, scene analysis, working memory, rule processing and emotional engagement (Patel, 2014). Alternatively, the additional demands imposed by musical listening in comparison to listening in general could lead to the acquisition of better general listening skills and different behavioural and neural mechanisms for listening. The correlation between auditory working memory and years of musical experience is parsimoniously explained by the acquisition of listening skills (Kraus et al., 2012; Lad et al., 2020). Different listening strategies are suggested by functional imaging of musicians carrying out listening tasks that suggest different neural systems for these in musicians (Gaab and Schlaug, 2003).

We favour an explanation for the specific correlation we have shown between musical sophistication and auditory working memory based on experience-dependent plasticity. The possible bases include changes in the ability of musicians to create and use mental stimulus representations. This would affect working-memory encoding and retrieval. Alternatively, training or experience might lead musicians to use auditory working memory resources more efficiently in the experimental task, which would affect retention. Further work could define specific differences in the behavioural components of tonal working memory in musicians, and in the brain network for this including auditory frontal and hippocampal cortex (Kumar et al., 2020, 2016).

## Materials and Methods

This study was pre-registered on the Open Science Framework online registry which can be found on https://osf.io/2n58c. Raw data is available online linked to the same registry and can be used in line with the CC0 1.0 Universal License.

### Participants

102 participants were recruited for two online experiments. In Experiment 1, 54 participants were identified from a local university database for behavioural experiments and in Experiment 2, 48 participants were recruited from prolific.com, an online website for identifying participants interested in behavioural experiments.

### Experimental Task

Participants performed the task via a web app. The interface played a video with instructions to each section of the task and participants were able to practice a trial from each stimulus modality of the working memory section once to familiarise themselves with the task. The entire task was divided into 3 sections (working memory task, questions from a questionnaire and non-verbal reasoning task) which were split further into subsections to reduce the monotony of performing one task over longer periods. Participants had an untimed break after 10 working memory trials, 6 questions or 4 non-verbal reasoning trials. Demographic information such as age and sex, and information regarding the set up used while performing the task with regards to the computer (desktop or laptop only) and headphones (in-ear or over-ear) were obtained after the experiment.

Experiment 1 was designed to test working memory precision in two different visual and auditory domains. For vision, colour hue stimuli from 0 to 300 and the rate of change of a black box to white with a modulation rate between 5 and 20 Hz were chosen. For audition, pure tones from 440 to 880 Hz and white noise modulated with a sine wave (100% depth) between 5 and 20 Hz were chosen. In Experiment 2, only pure tone and amplitude modulated auditory stimuli were used.

A schematic diagram of Experiment 1 is shown in Fig 1. Participants were asked to keep a visual or auditory stimulus in mind and ‘find’ the stimulus on a horizontal scale that they could interact with after a delay. A black cross at the centre of the screen with a white background marked the start of a trial. The initial stimulus was played for 1 sec, followed by a visual and auditory masker. The visual masker was designed as a 400 × 200 pixel screen filled with 10 × 10 pixel squares which randomly changed colour at a rate of 60 Hz. This was accompanied by the auditory masker which consisted of randomly generated 50 ms tone pips (from 440 to 880 Hz) for 0.5 seconds. After a delay of 2 to 4 seconds, participants viewed a 800 px horizontal line with a mouse-movable marker. To mark the beginning of the ‘Matching’ phase, a random probe stimulus (auditory or visual, depending on the trial) was played for 1 sec and an inverted red pointer was shown where *this* stimulus was located on the scale. Participants could freely move the marker and click to generate the stimulus at *the clicked* location for 1 sec. When they were satisfied that their click matched the original stimulus of interest, they could press the Return key on a keyboard. The parameter space for the stimulus of interest was mapped linearly to the pixel location of the horizontal scale. The extremes (10% most leftward and 10% most rightward) of the scale were not used for mapping as pilot studies indicated that performance is non-gaussian and skewed when stimuli are matched at these boundaries.

A schematic diagram of Experiment 2 is shown in Fig 3. In this experiment, along with the working memory task, participants performed a perceptual matching task interleaved in blocks. The latter began at the ‘Matching’ phase. For every trial participants were played an initial probe sound followed by the stimulus sound, after 100 ms, that needed to be matched. Participants had to make repeated match attempts to find the location on the horizontal scale where the two sounds matched exactly.

Participants completed the general Goldsmiths Musical Sophistication Index (Gold-MSI) questionnaire, containing 18 questions allowing for the scoring of a general musical sophistication factor (Müllensiefen et al., 2014). It is a self-report inventory that measures differences in skilled musical behaviours in the general or ‘non-specialist’ population. The questionnaire measures different factors associated with musical sophistication such as: active engagement, perceptual abilities, singing abilities and behaviours related to emotional responses to music. The maximum score is 156. Residuals from a linear regression with Age and Non-verbal reasoning scores (variables of no interest) were used for correlative analysis.

Participants also completed the Matrix Reasoning task from the 3rd Edition of the Wechsler Adult Intelligence Scale (Dugbartey et al., 1999). The original task has 30 questions arranged in order of difficulty. In order to increase engagement with the whole task, the last 12 even numbered questions were presented to participants. The first 3 questions were omitted as pilot studies revealed that these were seldom answered incorrectly. The total raw score was out of 12.

### Data Analysis

For every working memory trial, an error metric was calculated as the difference between the parameter of the initial stimulus and that which it was matched to. Each trial error was converted to a z-score for further analysis. For colour matching trials these errors were used to calculate working memory precision as the inverse of the standard deviation of the distribution of errors across colour trials of the whole experiment. For flash rate, frequency and amplitude modulation matching trials, the error was divided by the parameter of the stimulus to be matched first. This was carried out as for these trials the difference in percept of two images or sounds separated by the same parameter distance reduces as the value of the parameter increases i.e. two sounds pairs with a frequency of 440 and 480, and 840 and 880 are not perceived as similarly different and therefore the first pair has a greater ‘distance’ in musical notation. For Experiment 2, a similar analysis was carried out for auditory perception trials which yielded a metric for perceptual precision. In order to increase interpretability for precision values, these were converted into z-scores for individuals across all the participants.

Descriptive statistics and correlative analysis (Pearson’s Correlation and Linear Regression) was carried out using the Pingouin module in Python. Age and Matrix Reasoning scores were used as covariates in correlative analysis. Permutation analysis with replacement was used to generate 5000 samples for associations between frequency vs. AM rate working memory precision and adjusted Gold-MSI scores, and frequency working memory vs. perceptual precision and adjusted Gold-MSI scores. A *t*-test was subsequently used to test statistical significance for the difference in these associations. The Holm-Bonferroni method was used to test statistical significance after multiple comparisons.

## Acknowledgements

We would like to acknowledge all the participants in this study for their time and contribution.

